# HCMV promotes viral reactivation through the coordinated regulation of Notch signaling by UL8 and miR-UL36

**DOI:** 10.1101/2025.11.05.686718

**Authors:** Aaron Dirck, Nicole Diggins, Wilma Perez, Christopher Parkins, Michael Daily, Rebekah Turner, Luke Slind, Linh Nguyen, Daniel Malouli, Guanming Wu, Meaghan Hancock, Patrizia Caposio

## Abstract

Human cytomegalovirus (HCMV) establishes latency in CD34^+^ hematopoietic progenitor cells (HPCs), where reactivation is intimately linked to cellular differentiation. We demonstrate that the Notch signaling pathway, a key regulator of stem cell maintenance and differentiation, functions as a barrier to HCMV reactivation. Two viral gene products, UL8 and miR-UL36, modulate this pathway during reactivation. UL8 promotes degradation of the Notch3 receptor via the endosomal/lysosomal pathway, dependent on two tyrosine-based motifs (Y305/314) in its cytoplasmic tail. A UL8 mutant lacking these motifs fails to degrade Notch3, resulting in sustained Notch signaling and impaired reactivation *in vitro* and in humanized mice. Similarly, miR-UL36 reduces expression of Notch3 and the Notch transcription factor Recombination Signal Binding Protein For Immunoglobulin Kappa J Region (RBPJ), suppressing Notch signaling. Deletion of miR-UL36 inhibits reactivation, but this defect, like that of the UL8 mutant, can be rescued by pharmacologic Notch inhibition. Thus, HCMV employs multiple gene products to suppress Notch signaling and promote conditions conducive to reactivation. These findings reveal how HCMV manipulates host differentiation pathways to control latency and suggest therapeutic strategies to prevent viral recurrence in immunocompromised patients.

**Importance:** Human cytomegalovirus (HCMV) establishes lifelong latency, posing significant risks to transplant recipients and other immunocompromised individuals. Reactivation depends on progenitor cell differentiation, yet the viral mechanisms governing this process remain unclear. We identify Notch signaling as a major inhibitory pathway to reactivation and show that HCMV uses UL8 and miR-UL36 to suppress this pathway. UL8 degrades Notch3, while miR-UL36 downregulates Notch3 and RBPJ, together reducing Notch signaling and enabling reactivation. Mutant viruses lacking these regulators fail to reactivate efficiently, but this can be reversed by pharmacological inhibition of Notch. These findings establish Notch pathway suppression as a critical viral strategy for reactivation and highlight potential therapeutic targets for preventing HCMV disease.

## Introduction

Human cytomegalovirus (HCMV) remains a significant cause of morbidity and mortality in immunosuppressed individuals, such as those receiving solid organ or hematopoietic stem cell transplantations (1, 2). The virus persists latently in CD34⁺ hematopoietic progenitor cells (HPCs) and reactivates when immune surveillance is compromised, leading to systemic replication (3, 4). In stem cell transplant recipients, reactivation contributes to myelosuppression and graft failure, requiring treatment with antivirals such as Valganciclovir, which can exacerbate cytopenia and promote resistance (5). Despite the clinical burden, no vaccine exists and predicting reactivation remains difficult due to limited understanding of viral and host determinants.

Latency is defined by the presence of viral genomes without production of infectious virus, while reactivation involves overcoming intrinsic barriers to viral gene expression and the generation of new virions. This process is closely tied to HPC differentiation, as reactivation occurs when infected progenitors differentiate into macrophages or dendritic cells (6). Among signaling networks controlling hematopoietic fate, the Wnt/β-catenin and Notch pathways are particularly important. Wnt activation promotes differentiation (Wnt-ON), while Notch activation maintains stemness (Notch-ON) (7–11). Notch signaling involves cell-to-cell communication via the direct interaction of the Notch receptor on one cell with a membrane-bound ligand on a second cell (juxtacrine signaling). As such, Notch signals between cells in the bone marrow communicate fate decisions to hematopoietic cells. Mammals possess four different Notch receptors (Notch1-4), all of which have been detected on CD34^+^ HPCs. In mammalian signal-sending cells, members of the Delta-like (DLL1, DLL3, DLL4) and Jagged (JAG1, JAG2) families serve as ligands for Notch receptors. Upon ligand binding, the Notch extracellular domain (NECD) is cleaved away from the transmembrane-intracellular domain (TM-NICD) by TACE (TNF-α ADAM metalloprotease converting enzyme). The NECD remains ligand-bound and is internalized via ubiquitin-dependent endocytosis and recycled in the signal-sending cell. In the signal-receiving cell, γ-secretase releases the Notch intracellular domain (NICD) from the membrane, enabling its nuclear translocation, where it associates with the CSL (CBF1/RBPJ/Su(H)/Lag-1) transcription factor complex. This leads to activation of canonical Notch target genes such as HES and HEY family members, which function to block transcription of differentiation-related genes through recruitment of co-repressors including histone modification machinery (12, 13).

Previous studies implicate Notch signaling in various viral infections, including Epstein-Barr virus (EBV), Kaposi’s sarcoma-associated herpesvirus (KSHV), adenovirus, and human papillomavirus (HPV) (14–18). In EBV infection, EBNA2 mimics activated Notch, whereas in KSHV infection, LANA modulates Notch transcriptional responses (14, 15, 17, 18). While prior studies have shown that HCMV promotes abnormal differentiation in neural stem cells via modulation of Notch pathway members using exogenous expression of viral proteins such as pp71 and UL26 (19), the role of Notch signaling in HPCs, which have distinct developmental programs, has not yet been investigated.

Our prior work identified pUL7 as essential for reactivation through the Flt3 receptor, promoting monocyte differentiation (20). UL8, partially colinear with UL7, was shown to stabilize Wnt/β-catenin signaling by binding β-catenin and DVL2, promoting reactivation (21). Here, we uncover a complementary role for UL8 and miR-UL36 in suppressing Notch signaling. UL8 targets Notch3 for lysosomal degradation, while miR-UL36 downregulates Notch3 and RBPJ, together establishing a Wnt-ON/Notch-OFF environment that enables HCMV reactivation.

## Results

### Downregulation of Notch signaling stimulates HCMV replication

Our previous work uncovered the importance of Wnt/β-catenin signaling in regulating HCMV reactivation (21). Given that reactivation is tightly linked to the differentiation status of myeloid-lineage cells, and that Wnt and Notch pathways exert opposing effects on HPC stemness, with Wnt “ON” and Notch “OFF” favoring differentiation, we investigated the role of Notch signaling in HCMV infection of HPCs. Human embryonic stem cell (hESC)-derived CD34⁺ HPCs were infected with 2 TCID₅₀/cell of TB40/E-GFP for 48 hours. Viable CD34⁺GFP⁺ cells were sorted and seeded into long-term bone marrow culture (LTBMC) over stromal support to establish latency. After 12 days, cells were transferred onto permissive fibroblast monolayers in an extreme limiting dilution assay (ELDA) with cytokine-rich media to trigger differentiation and viral reactivation. Parallel cultures were mechanically lysed and plated to detect pre-formed virus, serving as a pre-reactivation control. To assess the role of Notch signaling in this model, we treated cultures with CB-103, a selective inhibitor of the CSL-NICD interaction that impairs the transcriptional response to Notch activation, either during latency or at the time of reactivation. Inhibition of Notch signaling during latency increased the amount of virus produced in latent cultures (Fig. 1A; pre-reactivation), suggesting impaired latency establishment and/or enhanced replication when Notch signaling is inhibited. CB-103 treatment only at the time of reactivation also enhanced the frequency of viral reactivation relative to controls (Fig. 1B), indicating that intact Notch signaling is inhibitory to virus replication at the time of reactivation. Importantly, CB-103 did not compromise CD34⁺ HPC viability or affect viral replication in fibroblasts (Fig. S1A–B). Additionally, we found that HCMV infection of CD34⁺ HPCs upregulated expression of the Notch target genes *HES1* and *HEY1* early after infection (Fig. 1C), without altering expression of Notch receptors (Fig. 1D), supporting the notion that Notch signaling is activated during latent infection and may contribute to latency establishment and/or maintenance. Together, these findings suggest that Notch signaling is important to limit virus replication during latency, and that downregulation of Notch signaling at the time of reactivation enhances the ability of HCMV to reactivate.

**Figure 1:**
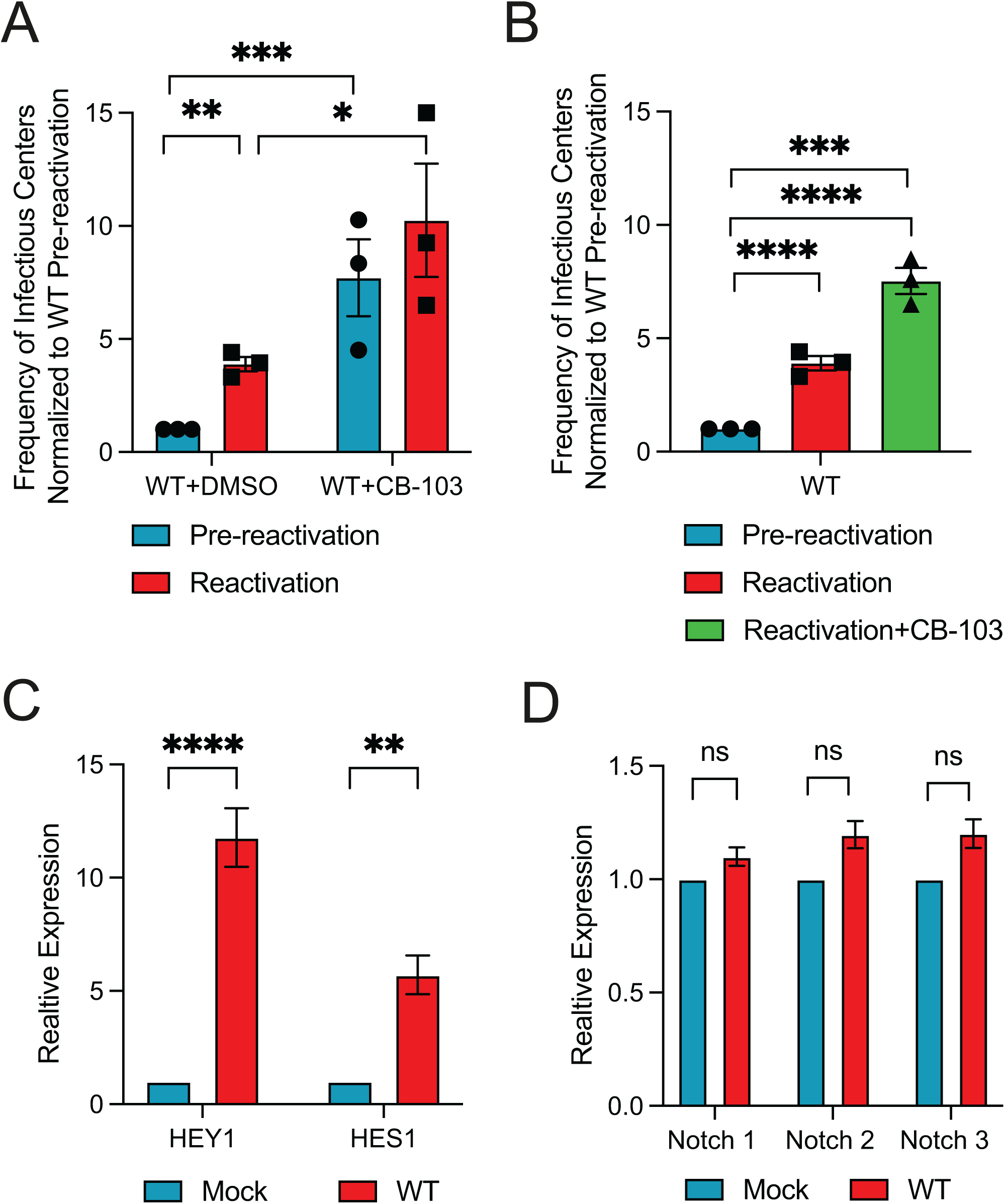
Inhibition of Notch signaling enhances HCMV replication. (A) Pure populations of CD34^+^ HPCs infected with WT HCMV were isolated by FACS at 48 hours post-infection and maintained in LTBMC medium in the presence of 10 μM of Notch inhibitor CB-103 or control DMSO, which was replenished at 7 days post-infection. At 14 days post-infection, viable CD34^+^ HPCs were seeded onto fibroblast monolayers plated in 96-well dishes by extreme limiting dilution in cytokine rich media for reactivation. An equivalent number of cells was mechanically disrupted and seeded in parallel to determine the infectious virus present in the cultures prior to reactivation (pre-reactivation). The frequency of infectious centers formation pre-and post-reactivation was determined 14 days later from the number of GFP^+^ wells at each dilution using extreme limiting dilution analysis. Data shown is the fold change in reactivation normalized to HCMV WT pre-reactivation. Bars represent the mean ± SEM of three independent experiments. Statistical significance was determined using two way-ANOVA with Tukey’s multiple comparison test (*p<0.05;**p<0.005;***p<0.0005). (B) Cells were infected and plated as in (A) except that 10 μM of Notch inhibitor CB-103 or DMSO was added to co-cultures only at the time of reactivation and replenished after one week. (***p<0.0005, ****p<0.0001). (C-D) CD34^+^ HPCs were mock infected or infected with WT HCMV for 48 hours, followed by sorting for GFP^+^, CD34^+^ viable cells. RNA was extracted from samples and subjected to qRT-PCR for the Notch target gene HEY1, HES1, and the receptors Notch1, 2, 3 (n=3; **p<0.01, ****p<0.0001 as determined by 2-tailed t test).

### UL8 interacts with components of the Notch pathway and promotes the degradation of Notch3

UL8 is a transmembrane protein with a highly glycosylated extracellular immunoglobulin (Ig)-like domain and a cytoplasmic tail characterized by two tyrosine-based endocytic motifs [YDRW (aa 305 to 308) and YGQV (aa 314 to 317)] and a PDZ-binding domain [DTEL (aa 321 to 324aa)], completely conserved among all strains (21, 22). In order to better understand the proteins and pathways associated with UL8, we employed an unbiased proximity-dependent labelling strategy using the promiscuous biotin ligase TurboID which biotinylates proteins within ∼10 nm of the tagged protein (23, 24). TurboID-mass spectrometry (MS) is advantageous compared to immunoprecipitation (IP)-MS because this system captures not only strong and direct protein-protein interactions, but also weak and/or transient interactions. Among biotin ligases, we used TurboID because it catalyzes proximity labelling with enhanced efficiency (24). We generated a fusion of TurboID to the C-terminus of UL8 in the TB40/E strain of HCMV and infected human fibroblasts for 96 hours in the presence of biotin. We have previously shown that TurboID does not alter pUL8 localization (21). Biotinylated proteins were isolated by streptavidin affinity purification (Fig. 2A) and analyzed by quantitative label-free mass spectrometry. Comparison of the MS hits in the UL8-TurboID samples to the wild-type virus control allowed us to identify significantly enriched host proteins in proximity to UL8. Hits from our proteomic analysis were refined by excluding proteins that were identified as likely contaminants based on comparison to the CRAPome database (25). Our revised list included 164 host proteins and five viral proteins (UL16, US24, US16, US15, UL130) (Dataset S1).

**Figure 2.**
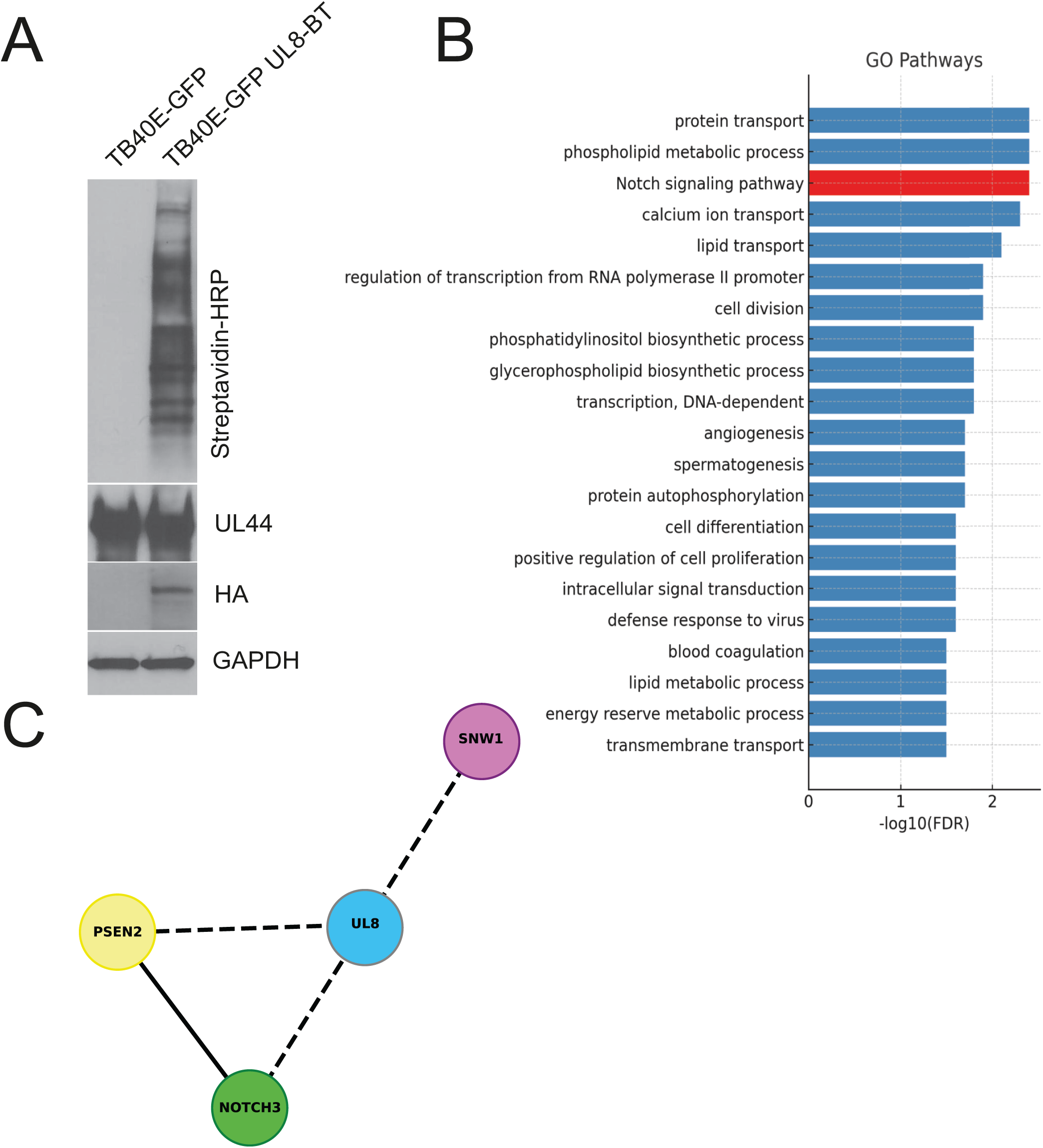
HCMV UL8 interacts with components of the Notch signaling pathway. (A) NHDF were infected with TB40/E-GFP-UL8-BT or TB40/E-GFP at an MOI of 2. At 3 days post-infection, the cells culture medium was supplemented with biotin (50 μg/ml) for 6 hours. Labeling and infection were confirmed by immunoblot on whole cell lysates using the indicated primary antibodies. (B) Gene Ontology (GO) enrichment analysis of proteomic data identifies significantly enriched biological processes. The bar plot shows the top GO terms ranked by statistical significance (-log₁₀(FDR)). (C) Network diagram (https://cytoscape.org) illustrating the proximity-based associations of the HCMV protein UL8 (grey) with key components of the Notch signaling pathway (colored nodes: AGO3, NOTCH3, PSEN2, SNW1, TNRC6A, and TNRC6C). Dashed lined indicate interactions identified by BioID-MS analysis and solid lines represent protein-protein interactions derived from the STRING database (https://string-db.org).

Gene Ontology (GO) enrichment analysis of these UL8-associated proteins revealed a range of significantly enriched biological processes, including protein transport, phospholipid metabolism, and calcium ion transport. Among these, the Notch signaling pathway emerged as one of the most significantly enriched terms (Fig. 2B). Focusing specifically on the Notch signaling pathway, we constructed a network diagram highlighting the proximity-based associations between UL8 and key Notch pathway components (Fig. 2C). The analysis identified three Notch-related proteins, NOTCH3, Presenilin 2 (PSEN2), and SNW domain-contain protein 1 (SNW1), in close proximity to UL8 (FDR<0.1, Dataset S1). Among these, Notch3 was particularly noteworthy. Notch3 is a canonical receptor in the Notch signaling pathway, playing a crucial role in cell differentiation and immune regulation (26). In our UL8-associated proximity network, Notch3 emerged as a central hub, interacting with multiple other Notch pathway components. This hub-like position suggests that Notch3 may serve as a critical mediator of UL8’s function during HCMV infection. This finding forms the basis for our subsequent mechanistic studies to explore the UL8–Notch3 axis in HCMV-infected fibroblasts.

First, we aimed to validate the interaction between UL8 and Notch3 and to identify the specific residues mediating this interaction. We focused on the tyrosine motifs at position 305 and 314 based on previous studies demonstrating their importance for UL8 internalization (22). Given that endocytic trafficking often facilitates receptor engagement, we hypothesized that these motifs may also contribute to UL8’s ability to bind the Notch receptor. To this end, we generated adenoviral constructs expressing wild-type UL8, a UL8 mutant carrying tyrosine-to-alanine substitutions at position 305 and 314 (UL8 Y305A; UL8 Y314A) and the double mutant UL8 Y305/314A. As controls, we included Ad-UL7 and a mutant UL8 lacking the DTEL motif (Ad-UL8 DTEL/AAAA, referred to as mutDTEL), since Notch3 does not contain a PDZ domain and therefore would not be expected to interact via this motif (Fig 3A). Fibroblasts were transduced with the adenoviral constructs for 48 hours, followed by a pull-down assay using a Notch3-specific antibody. As shown in Figure 3B, UL8 interacts with Notch3, and efficient interaction depends on the on the tyrosine-based endocytic motifs Y305 and 314. Additionally, when we performed western blot analysis on total protein lysates prior to immunoprecipitation, we observed that Notch3 levels were decreased in the presence of UL8 and UL8 mutDTEL, suggesting that UL8 expression promotes Notch3 degradation independently of its PDZ-binding motif (Fig. 3B). Previous work by Carmon-Perez *et al*. demonstrated that UL8 undergoes degradation via the endosomal/lysosomal pathway (22). Given the interaction between UL8 and Notch3, we hypothesized that UL8 may mediate Notch3 degradation by co-trafficking to the endo-lysosomal compartment. To test this hypothesis, HEK293 cells were co-transfected with plasmids expressing UL8 and the active intracellular Notch3 domain (NICD), in the presence or absence of Leupeptin and Brefeldin-A1, two well-characterized inhibitors of endosomal trafficking. As shown in Figure 3C, treatment with Leupeptin and Brefaldin-A1 rescued the degradation of both Notch3 and UL8, consistent with previously published findings showing that these inhibitors stabilize UL8. These results support our hypothesis that UL8 and Notch3 are co-trafficked and degraded through a shared endo-lysosomal pathway.

**Figure 3:**
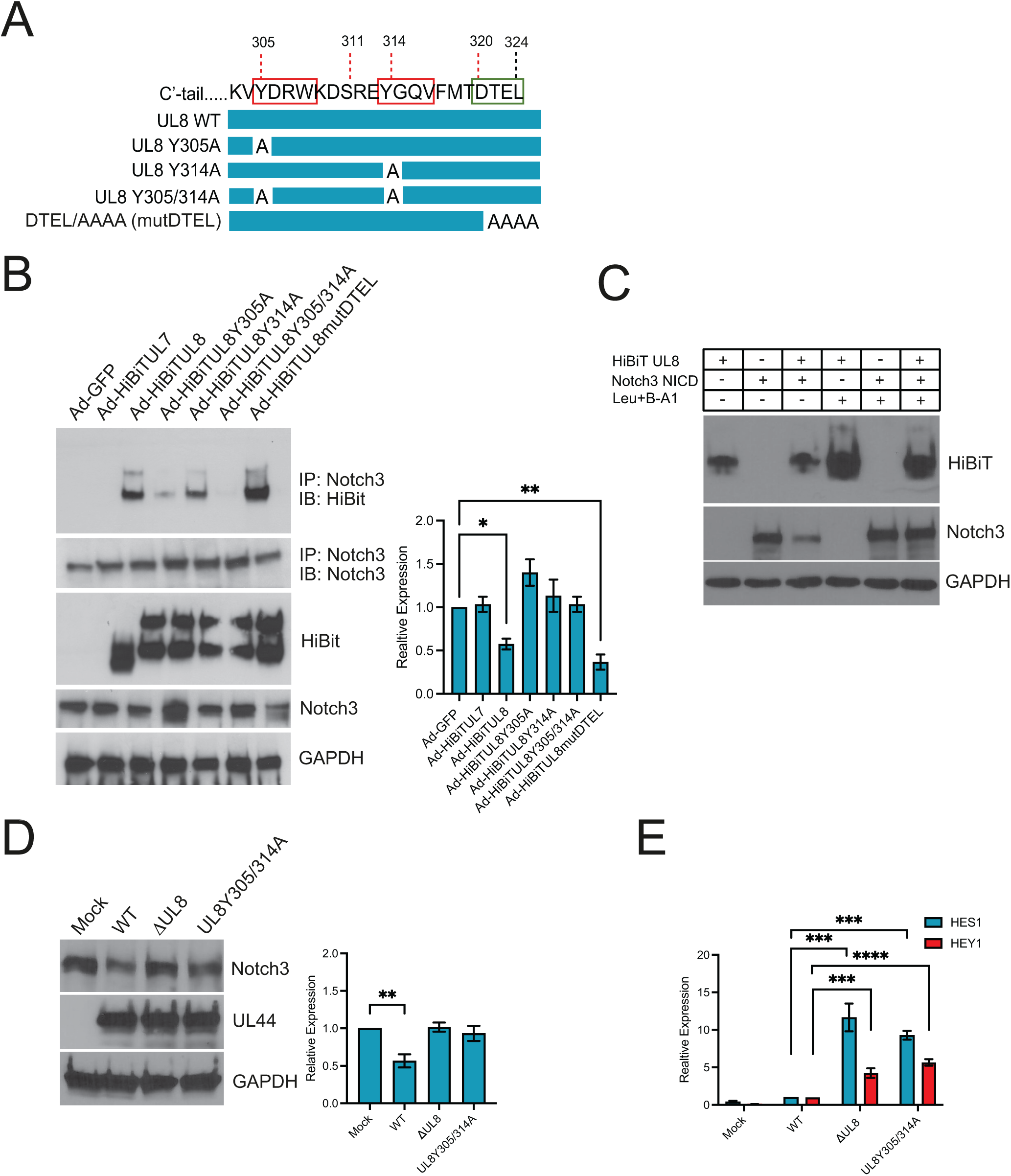
UL8 associates with Notch 3 and promotes its degradation. (A) Schematic of the UL8 C-terminal region showing the introduced mutations in the tyrosine binding motifs at Y305 and Y314, as well as in the PDZ-binding domain (DTEL). (B) NHDF were transduced with adenoviral constructs expressing GFP, UL7, UL8, UL8Y305A, UL88Y314A, UL8 Y305/314A, or UL8-mutDTEL for 48 hours. Using rabbit anti-Notch3 antibody, UL8 was immunoprecipitated from cell lysates, and proteins separated by SDS-PAGE. Immunoprecipitated proteins were detected by Western blotting using the NanoGlo HiBiT Blotting system. A Western blot of total lysate is shown with GAPDH as a loading control. Relative protein expression of three independent experiments was quantified using Image J. Values are means±standard error of the means (SEM) (error bars). Statistical significance was determined by unpaired Student’s t-test. *\*, p<0.05; **, p<0.005* compared to Ad-GFP. (C) HEK293 cells were co-transfected with plasmids expressing UL8-V5 and Notch3 NICD. After 24 h, cells were left untreated or incubated with 50 μM leupeptin (Leu) and 20 nM bafilomycin A1 (B-A1) for 24 hours. Cells were then lysed, and collected samples were subjected to SDS-PAGE and immunoblotting using anti-V5, and anti-Notch3 antibodies. GAPDH was used as a loading control. A representative Western blot, out of three, is shown. (D) NDHF were infected with HCMV TB40/E WT, ΔUL8, or UL8 Y305/314A at an MOI of 1. At 96 hours post-infection protein lysates were generated and immunoblotted for Notch3, HCMV UL44 and GAPDH as a loading control. Densitometry was performed using ImageJ software for 3 independent experiments and plotted in right-hand panels. Values are means ±standard error of the means (SEM) (error bars). Statistical significance was determined by unpaired Student’s t-test. \**\*, p<0.005* compared to Mock-infected cells. (E) RNA was harvested from cells infected as in (D) and subjected to qRT-PCR using primers for the Notch target genes HES1 and HEY1 (n=3; ***p<0.001, ****p<0.0001 by 2 tailed t test).

Finally, we sought to assess the impact of UL8 on Notch3 protein levels and downstream Notch target genes expression during lytic HCMV infection. Fibroblasts we infected with wild-type HCMV, a UL8 deletion mutant (ΔUL8), or a UL8 mutant carrying tyrosine-to-alanine substitutions at position 305 and 314 (UL8Y305/314A). As shown in Figure 3D, Notch3 protein levels were reduced in cells infected with wild-type virus compared to those infected with the ΔUL8 or UL8 Y305/314A mutants, indicating that UL8 promotes Notch3 degradation during lytic infection. Consistently, transcript levels of the Notch3 target genes *HES1* and *HEY1* were elevated at 96 hours post-infection in cells infected with ΔUL8 and UL8 Y305/314A mutants relative to wild-type infected cells, supporting the hypothesis that UL8-mediated Notch3 degradation suppresses downstream transcriptional activity (Fig. 3E). Together, these findings uncover a novel mechanism by which UL8 modulates host cell signaling and establishes UL8 as a previously unrecognized regulator of Notch3 stability and activity during HCMV infection.

### miR-UL36 disrupts Notch signaling during lytic infection

Previous studies have implicated HCMV-encoded miRNAs in regulating signaling pathways necessary for latency establishment and reactivation (27–31). Thus, we hypothesized that HCMV miRNAs may contribute to regulating Notch signaling during infection. To test this hypothesis, we screened HCMV miRNAs for their ability to reduce signaling in a luciferase-based Notch signaling assay. HEK293 cells were transfected with a plasmid expressing a luciferase reporter driven by a high affinity 4X CSL binding site and a plasmid expressing a constitutively active NICD along with negative control or HCMV miRNA mimics. As shown in Figure 4A, we noted that miR-UL36 caused a reduction in luciferase expression not observed with miR-US33 or miR-UL112-3p miRNA mimics, suggesting that miR-UL36 interferes with signaling mediated by the Notch intracellular domain.

**Figure 4:**
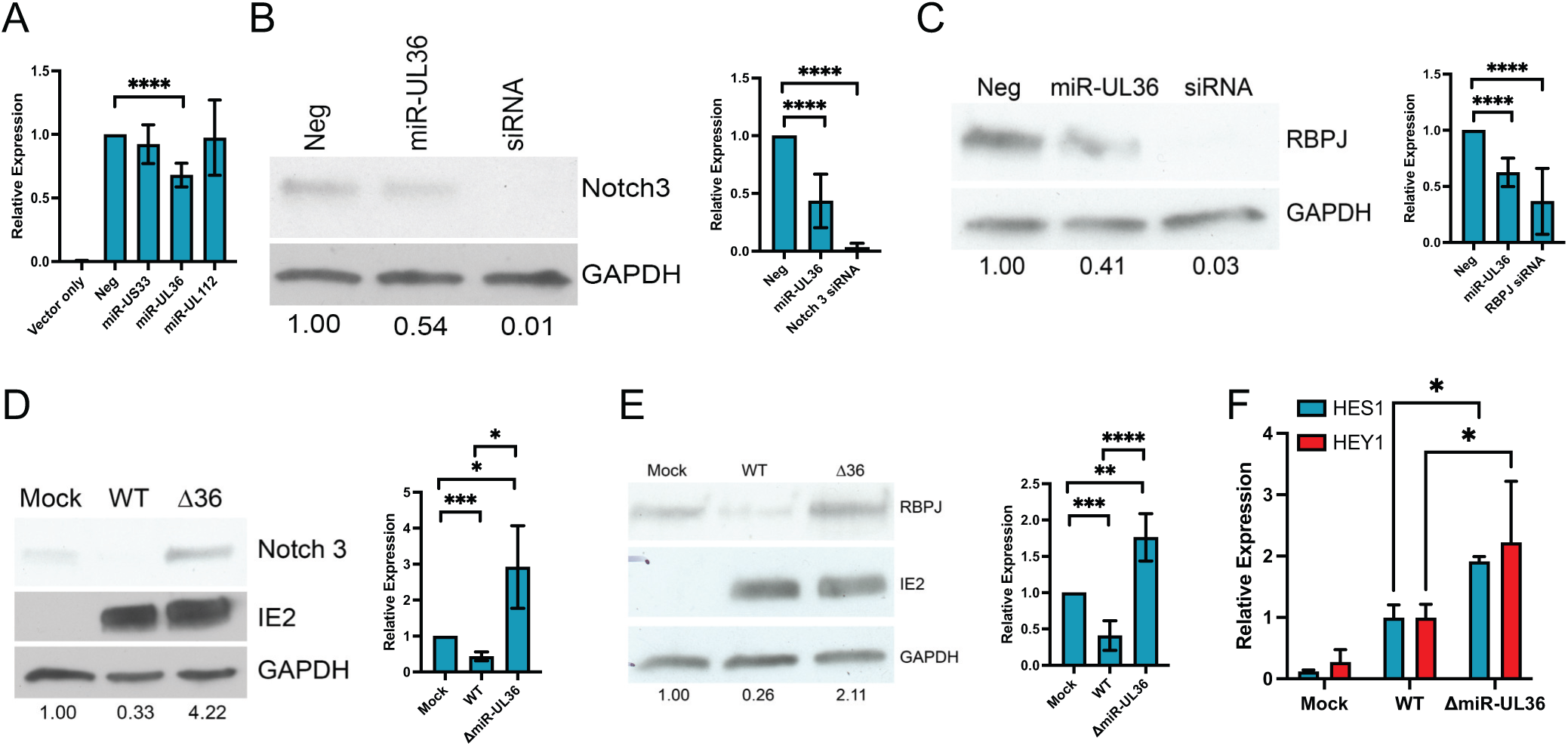
miR-UL36 impairs Notch signaling through targeting Notch3 and RBPJ. **(A)** HEK293 cells were transfected with a 4XCSL-Luciferase construct as well as a plasmid expressing the intracellular domain of Notch3 and either negative control miRNA mimic, miR-US33, miR-UL36 or miR-UL112. Protein lysates were harvested 24 hours post-transfection and luciferase expression was monitored using a luminometer. (n=3; p<0.05 calculated using a one sample t and Wilcoxon test). **(B-C)** NHDF were transfected with negative control miRNA mimic, miR-UL36 or siRNA against Notch3 (B) or RBPJ (C). Protein lysates were harvested 48 hours post-transfection and immunoblots were performed using the indicated antibodies. Densitometry was performed using ImageJ software for 3 independent experiments and plotted in the right-hand panels. **(D-E)**. NHDF were mock-infected or infected with WT HCMV or a miR-UL36 mutant virus for 48 hours. Protein lysates were harvested, and immunoblotting was performed using the indicated antibodies. Densitometry was performed using ImageJ software for 3 independent experiments and plotted in the right-hand panels. *p<0.05, **p<0.01, ***p<0.001, ****p<0.0001 as determined by 2-tailed t test. **(F)** RNA was harvested from cells infected as in (D-E) and subjected to qRT-PCR using primers for the Notch target genes HES1 and HEY1. (n=3; *p<0.05 by 2 tailed t test).

We further investigated the ability of miR-UL36 to regulate expression of two key components of the Notch signaling pathway, the receptor Notch3 and the transcriptional regulator RBPJ. As shown in Figure 4B and C, expression of a miR-UL36 miRNA mimic in HEK293 cells reduced both Notch 3 and RBPJ protein levels by ∼50%. Next, we generated a miR-UL36 mutant virus whereby the miR-UL36 sequence, encoded within the intron of the UL36 transcript, was deleted. pUL36 is an inhibitor of apoptosis and necroptosis and plays a role in regulating innate immune signaling (32, 33), thus we wanted to ensure that mutation of miR-UL36 had no effect on pUL36 expression or virus replication. To assess pUL36 expression we generated a recombinant virus where the FLAG sequence was added to the C terminus of UL36 and additionally generated a recombinant virus where the miR-UL36 sequence was mutated. We infected human fibroblasts with WT, UL36FLAG and ΔmiR-UL36-UL36FLAG viruses and harvested protein for immunoblot. As shown in Figure S2A, we did not observe any notable difference in UL36 protein accumulation in the absence of miR-UL36 expression. We also assessed growth kinetics of the ΔmiR-UL36 virus compared to WT following infection of fibroblasts (Fig. S2B) and did not note any differences in growth when miR-UL36 was mutated. Infection of fibroblasts with the ΔmiR-UL36 virus resulted in significantly higher levels of Notch 3 (Fig. 4D) and RBPJ proteins (Fig. 4E) compared to WT-infected cells. To investigate whether the enhanced expression of Notch signaling components that occurs in the absence of miR-UL36 results in enhanced Notch signaling, we measured HES1 and HEY1 transcript levels in fibroblasts mock-infected or infected with WT or ΔmiR-UL36 viruses. As shown in Figure 4F, we observed a significant increase in HES1 and HEY1 transcripts in the absence of miR-UL36, compared to WT-infected cells, indicating that miR-UL36, like UL8, blocks Notch-mediated transcriptional responses during lytic infection of fibroblasts.

### UL8-mediated degradation of Notch 3 is essential for HCMV reactivation *in vitro*

To assess the functional relevance of the UL8-Notch3 interaction in CD34^+^ HPCs, we measured expression levels of the Notch target gene *HES1* at 48h post-infection, a time when UL8 is expressed in CD34^+^ HPCs prior to the establishment of latency (34). As shown in Figure 5A, CD34^+^ HPCs infected with the UL8 Y305/314A mutant exhibited elevated *HES1* expression compared to those infected with wild-type virus, consistent with the observations in infected fibroblasts and uncovering a role for UL8 in suppressing Notch signaling in hematopoietic cells.

**Figure 5:**
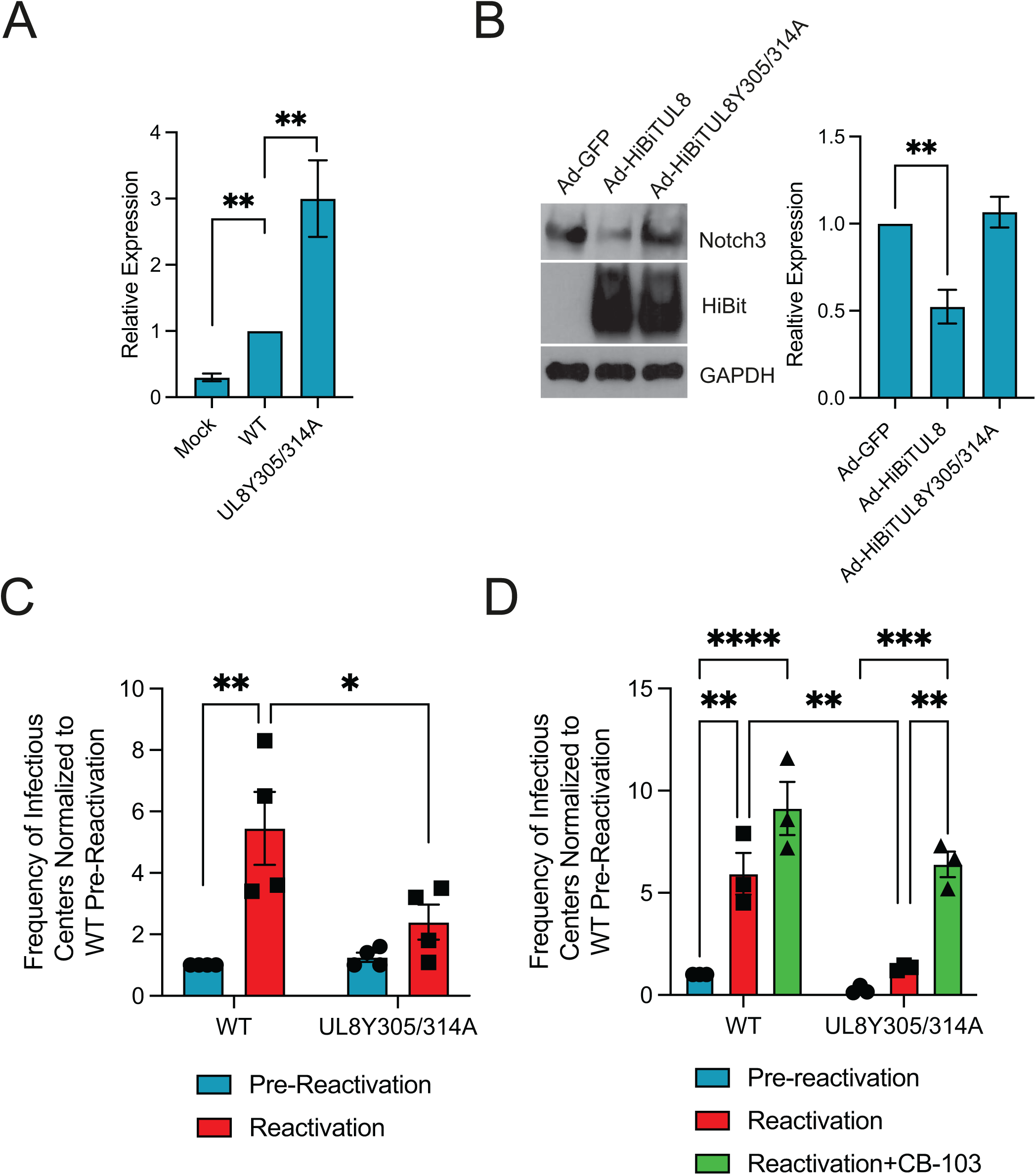
UL8 modulation of Notch signaling is required for viral reactivation in CD34^+^ HPCs. **(A)** CD34^+^ HPCs were mock infected or infected with WT HCMV or UL8 Y305/314A virus for 48 hours, followed by sorting for GFP^+^, CD34^+^ viable cells. RNA was extracted from samples and subjected to qRT-PCR for the Notch target gene HES1. (n=3;**p<0.001, by 2-tailed t test). **(B)** CD34^+^ HPCs were transduced for 48 hours with adenoviruses expressing GFP, UL8 and UL8 Y305/314A. After sorting for GFP^+^, CD34^+^ viable cells, proteins were extracted, and immunoblotting using the indicated antibodies was performed. Densitometry was performed using ImageJ software for 3 independent experiments and plotted in the right-hand panels. Values are means ±standard error of the means (SEM) (error bars). Statistical significance was determined by unpaired Student’s t-test. **p<0.005, compared to Ad-GFP. **(C)** CD34^+^ HPCs were infected with WT or UL8 Y305/314A viruses and latency and reactivation experiments were performed as in Figure 1. **(D)** CD34^+^ HPCs were infected with WT or UL8 Y305/314A viruses and latency and reactivation experiments were performed as in Figure 1. 10 μM of Notch inhibitor CB-103 was added to co-cultures at the time of reactivation and replenished after one week.

To further investigate the role of UL8 in regulating Notch signaling in HPCs, we infected CD34^+^ HPCs with adenoviruses expressing either GFP (negative control), wild-type UL8, or the UL8 Y305/314A mutant. 48 hours post-infection infection, GFP-positive cells were sorted by FACS, and protein lysates were prepared. Immunoblotting with antibodies against Notch3 revelated that UL8 expression markedly reduced Notch3 protein levels compared to Ad-GFP-infected cells (Fig. 5B). In contrast, expression of the UL8 Y405/314 mutant did not significantly alter Notch3 levels relative to Ad-GFP control, suggesting that UL8 mediates Notch3 downregulation outside the context of HCMV infection, and that the tyrosine motifs at position 305 and 314 are critical for UL8-mediated Notch3 downregulation during infection of HPCs.

We have previously demonstrated that deletion of UL8 impairs viral reactivation (21), although the contribution of regulating Notch signaling to this phenotype is unknown. Therefore, we performed our *in vitro* latency and reactivation assay as previously described using the UL8Y305/314A mutant. As shown in Figure 5C, mutation of the UL8 tyrosine based-endocytic motifs blocked viral reactivation, supporting the hypothesis that lysosomal degradation of Notch3 is an essential function of UL8 during reactivation. Quantitative analysis of HCMV genomes in cells infected with either wild-type or UL8 Y305/314A mutant virus at 14 days post-infection (dpi) revealed comparable DNA levels (Fig. S3A), confirming that the observed reactivation defect was not due to loss of viral genomes during the latency period. Furthermore, no replication defect was noted upon multi-step growth curve analysis (Fig. S3B). Finally, we sought to determine whether inhibition of Notch signaling with CB-103 could rescue the reactivation defect of the UL8 Y305/314A mutant virus. As shown in Figure 5D, addition of CB-103 at the time of reactivation partially restored the ability of the mutant virus to reactivate, further supporting the role of UL8 in modulating Notch signaling during reactivation. Together, these results demonstrate that UL8-mediated degradation of Notch3 is a critical step for efficient HCMV reactivation in CD34^+^ HPCs.

### miR-UL36 regulation of Notch signaling is necessary for efficient reactivation from latency in CD34^+^ HPCs

In order to determine whether miR-UL36 also regulates Notch signaling in hematopoietic cells, we assessed HES1 levels in CD34^+^ HPCs at 2 days post-infection using WT or ΔmiR-UL36 viruses. As shown in Figure 6A, viral infection induced HES1 expression compared to mock-infected cells, while infection with the ΔmiR-UL36 virus resulted in significantly increased HES1 transcripts, suggesting that miR-UL36 indeed regulates Notch signaling during infection of CD34^+^ HPCs. Next, we infected CD34^+^ HPCs with adenoviruses expressing either the *C. elegans* miRNA miR-67 as a negative control or expressing miR-UL36. 48 hours after infection, cells were sorted by FACs using CD34 and GFP expression and protein lysates were generated from Ad-miR-67 and Ad-miR-UL36 transduced cells. As shown in Figure 6B, expression of miR-UL36 reduced Notch3 and RBPJ protein levels in infected CD34^+^ HPCs compared to Ad-miR-67 infected cells, further supporting a role for miR-UL36 in regulating Notch signaling during infection of HPCs.

**Figure 6:**
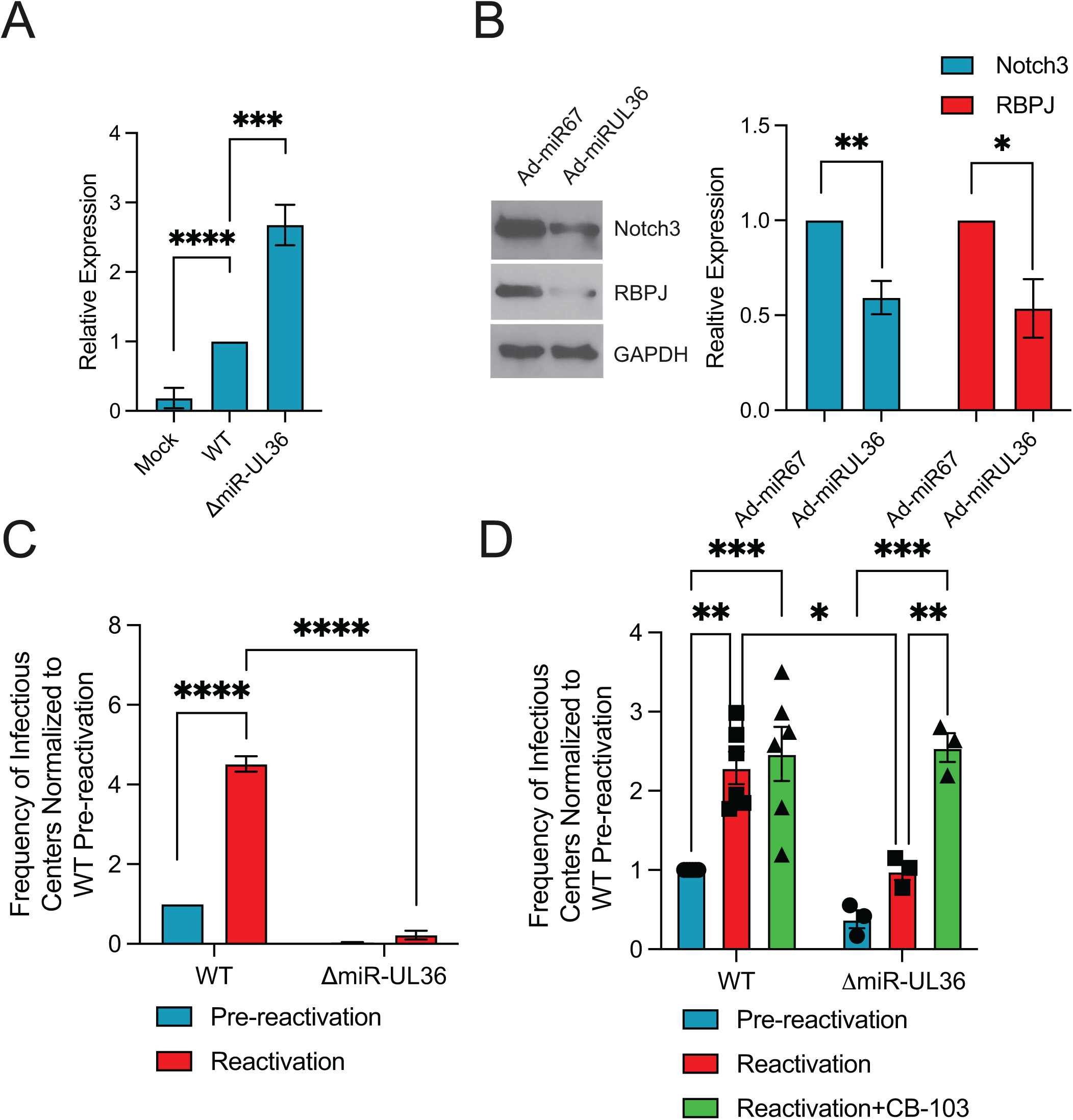
miR-UL36 regulates Notch signaling during infection of CD34^+^ HPCs. **(A)** CD34^+^ HPCs were mock infected or infected with WT HCMV or ΔmiR-UL36 virus for 48 hours, followed by sorting for GFP^+^, CD34^+^ viable cells. RNA was extracted from samples and subjected to qRT-PCR for the Notch target gene HES1. (n=3; ***p<0.001, ****p<0.0001, by 2-tailed t test). **(B)** Adenoviruses encoding GFP and expressing *C. elegans* miR-67 as a negative control, or miR-UL36 were used to infect CD34^+^ HPCs. After sorting for GFP^+^, CD34^+^ viable cells, proteins were extracted and immunoblotting using the indicated antibodies was performed. Densitometry was performed using ImageJ software for 3 independent experiments and plotted in the right-hand panels. Values are means ±standard error of the means (SEM) (error bars). Statistical significance was determined by unpaired Student’s t-test. **(C)** CD34^+^ HPCs were infected with WT or ΔmiR-UL36 viruses and latency and reactivation experiments were performed as in Figure 1. **(D)** CD34^+^ HPCs were infected with WT or ΔmiR-UL36 viruses and latency and reactivation experiments were performed as in Figure 1. 10 μM of Notch inhibitor CB-103 was added to co-cultures at the time of reactivation and replenished after one week.

To assess the importance of miR-UL36 and its regulation of Notch signaling in latency and/or reactivation in CD34^+^ HPCs we infected cells with WT or ΔmiR-UL36 viruses and performed reactivation assays as described above. As shown in Figure 6C, a lack of miR-UL36 expression results in a significant defect in the ability to produce new virus after reactivation stimulus. Furthermore, we detected no difference in viral genome levels when comparing WT and ΔmiR-UL36-infected cells (Fig. S3C), nor replication of the viruses in multi-step growth curves (Fig. S3D).

Finally, to determine whether regulation of Notch signaling was one of the functions of miR-UL36 required at the time of reactivation, we treated WT and ΔmiR-UL36 infected cells with DMSO or the Notch inhibitor CB-103 at the time of reactivation and replenished the drugs again after one week. As shown in Figure 6D, we observed that treatment with CB-103 enhanced the reactivation of WT virus, similar to Figure 1B. Furthermore, CB-103 treatment also significantly enhanced the reactivation of the ΔmiR-UL36 virus, indicating that regulation of Notch signaling is a major function of miR-UL36 at the time of reactivation in CD34^+^ HPCs.

### UL8 and miR-UL36 are important for reactivation in a humanized mouse model of latency

Given the importance of UL8 and miR-UL36 in reactivation from latency *in vitro*, we next asked whether viral regulation of Notch signaling plays a critical role in HCMV reactivation from latency *in vivo.* Using our humanized mouse model, human CD34^+^ HPCs were engrafted into NSG mice followed by injected with HCMV-infected fibroblasts. Four weeks later, mice were treated with G-CSF and AMD3100 to induce mobilization of myeloid cells from the bone marrow, thereby promoting viral reactivation (35). In mice infected with wild-type HCMV, the G-CSF/AM3100 mobilization triggered a marked increase in viral DNA copy number in the spleen and liver (Fig. 7A-B), consistent with efficient reactivation. In contrast, mice infected with viral mutants either containing the Y305/314A mutation in UL8 or lacking miR-UL36, both of which were shown *in vitro* to modulate Notch signaling (Figs. 5C, 6C), failed to exhibit a significant increase in viral DNA in either tissue following mobilization. These *in vivo* findings closely recapitulate our *in vitro* results and strongly suggest that UL8-and miR-UL36-mediated regulation of the Notch pathway is also essential for HCMV reactivation from latency in humanized mice.

**Figure 7.**
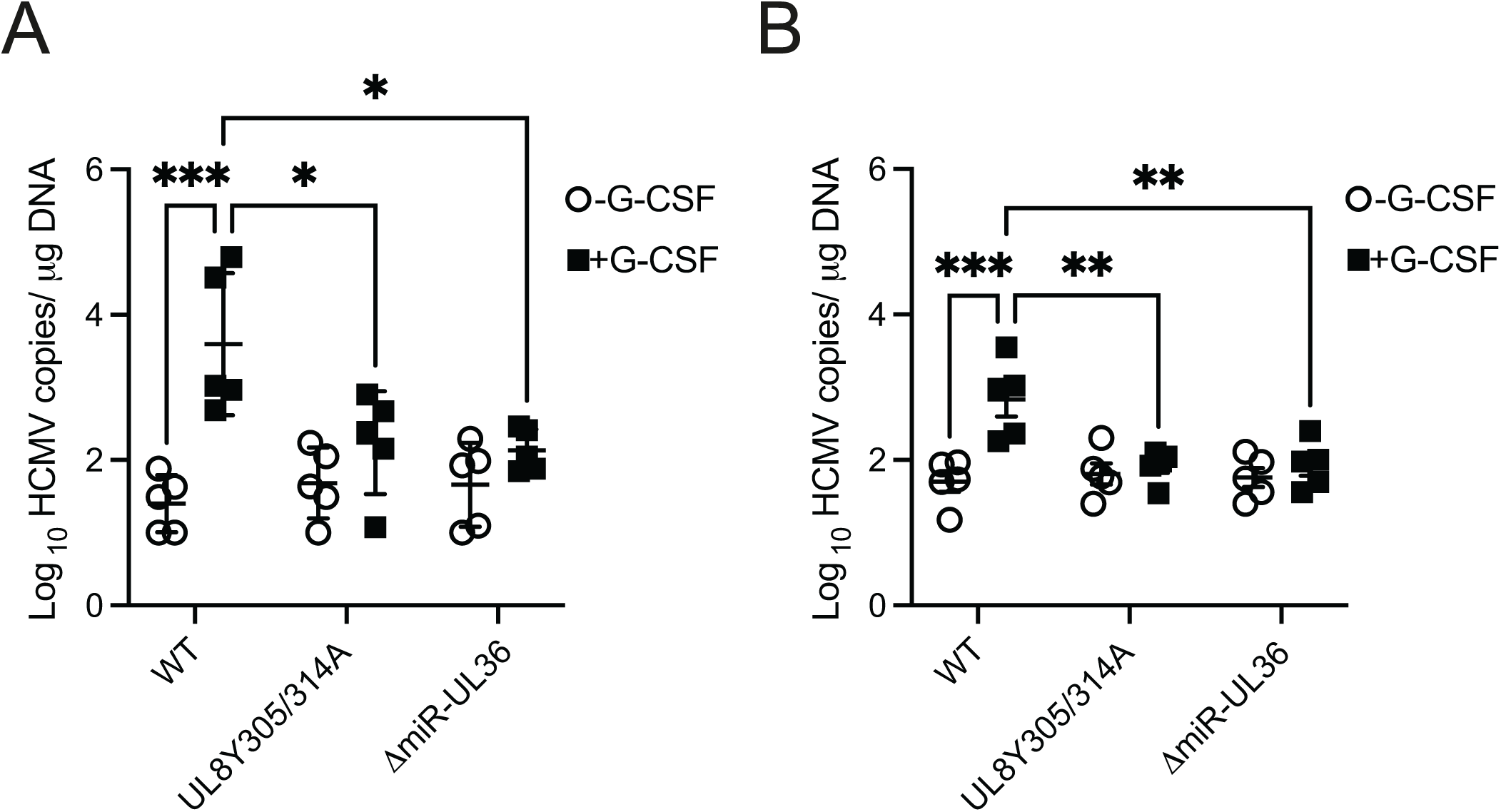
**UL8 and miR-UL36 are required for efficient viral reactivation *in vivo.*** Sublethally irradiated NOD-*scid* IL2Rγcnull mice were engrafted with human CD34^+^ HPCs (huNSG) and subsequently injected with human fibroblasts previously infected with HCMV WT, UL8 Y305/314A, or ΔmiR-UL36 viruses. At 4 weeks post-infection, viral reactivation was triggered by treating latently-infected (n=5) animals with G-CSF and AMD-3100 as previously described (35). At 1 week post-treatment, treated and untreated mice were euthanized and tissues harvested. Total genomic DNA was isolated from spleen tissue (A) or liver tissues (B), and HCMV genomes were quantified using quantitative PCR with primers and probe specific for the *UL141* gene. Statistical significance was determined using two-way ANOVA, followed by Bonferroni’s multiple comparison test (*\*, p<0.01; **, p<0.001; ***p<0.0001*).

## Discussion

The work presented herein supports a critical role for viral regulation of Notch signaling in reactivation from latency in CD34⁺ HPCs. Using a small molecule inhibitor, we show that inhibition of Notch signaling during latent infection or at the time of reactivation enhances virus replication in HPCs (Fig. 1A, B). Thus, downregulation of Notch signaling appears to be a necessary step for virus replication. We identify two viral gene products, the early-late protein UL8 and the HCMV-encoded miRNA miR-UL36, as key regulators that stimulate replication through suppression of Notch signaling. UL8 interacts with Notch3 via tyrosine residues in its C-terminal tail, promoting its degradation (Fig. 3), while miR-UL36 reduces Notch3 and the transcriptional regulator RBPJ protein levels (Fig. 4). Together, they limit Notch-dependent transcription in fibroblasts and CD34⁺ HPCs (Figs. 5A–B, 6A–B). Both UL8 and miR-UL36 are required for reactivation *in vitro* and *in vivo* (Figs. 5C, 6C, 7C–D), and Notch inhibitors can compensate for their loss (Figs. 5D, 6D), demonstrating that suppression of Notch signaling is essential for efficient reactivation. Given Notch’s role in maintaining HPC stemness and the link between reactivation and myeloid differentiation, we propose that UL8-and miR-UL36-mediated inhibition of Notch signaling promotes viral replication.

Early after infection, Notch target genes are transiently activated without changes in receptor expression (Fig. 1C, D). Activation of Notch signaling typically requires receptor-ligand engagement and nuclear translocation of NICD. Whether in the case of HCMV infection this activation reflects a host response or direct viral stimulation of the pathway remains unclear. We hypothesize that the functional result of activated Notch signaling is limiting HPC differentiation in order to prevent viral replication.

Thus, activation of Notch signaling early after infection of HPCs may help to reinforce the restrictive cellular environment necessary for latency establishment.

Although several viruses modulate the Notch pathway (14–18), direct interaction with the receptor is rare. The only previous example is the Human T-cell Leukemia Virus type 1 (HTLV-1) Tax protein, which prolongs the half-life of the Notch intracellular domain (NICD) thereby interfering with downstream transcriptional activity (7, 9, 10, 13, 26). Herpes Simplex Virus type 1 (HSV-1) encodes ICP0 which targets transcriptional co-activators for degradation rather than the receptor itself (20, 29, 36). Similarly, KSHV proteins such as Latency-associated nuclear antigen (LANA) and viral Interferon Regulatory Factors (vIRFs) suppress Notch pathway activity, with some evidence suggesting downregulation of Notch1, though not through direct receptor degradation (14). Our study identifies UL8 as a previously unrecognized modulator of the Notch signaling pathway, with a specific role in promoting Notch3 degradation during HCMV infection (Fig. 3D). Proximity-dependent biotin labeling revealed enrichment of Notch components in the UL8 interactome, with Notch3 as a central hub (Fig. 2B–C). This finding is particularly significant given the established role of Notch signaling in regulating cell differentiation, immune responses, and tissues homeostasis (7, 9, 10, 13, 26). Notch signaling serves as a critical regulator of progenitor cell fate decisions, balancing self-renewal with lineage commitment. High Notch activity maintains stemness and directs early T-cell specification, while its attenuation permits alternative differentiation programs, including B-cell and myeloid lineages (9, 10, 37). Within CD34⁺ hematopoietic progenitor cells, Notch– ligand interactions provided by the bone marrow niche act as checkpoints that influence whether cells persist in an undifferentiated state or undergo lineage progression (38). Thus, viral manipulation of Notch, particularly through direct degradation of receptors such as Notch3, has the potential to profoundly alter hematopoietic cell dynamics, skew immune lineage development, and create a cellular environment favourable to HCMV reactivation. The ability of HCMV to alter HPC differentiation and lineage development has been previously described in both *in vitro* and *in vivo* models (20, 29, 36) and our data support the notion that HCMV regulation of Notch signaling is one component of these processes.

Mechanistically, we show that UL8 binds Notch3 through two conserved tyrosine-based motifs (Fig. 3B), mediating co-trafficking to the endo-lysosomal compartment. Lysosomal inhibition stabilizes both UL8 and Notch3 (Fig. 3C), confirming degradation through this pathway. This activity is independent of UL8’s PDZ-binding motif, which mediates β-catenin interaction, indicating that UL8 controls Wnt-ON and Notch-OFF states via distinct domains. Functionally, UL8-mediated Notch3 degradation correlates with reduced HES1 and HEY1 expression (Fig. 3E), linking receptor turnover to transcriptional attenuation.

Previous work has uncovered essential roles for viral miRNAs in regulating signal transduction through numerous cell signaling pathways (39). Here we add regulation of Notch signaling via miR-UL36 targeting key pathway component Notch3 and RBPJ (Figs. 4, 6). Surprisingly, no canonical binding sites for miR-UL36 were found in Notch3 or RBPJ 3’UTRs, thus regulation of these proteins may be non-canonical or indirect. RBPJ can act as an activator or repressor and has been shown to limit CMV IE gene expression *in vivo* (40). Thus, miR-UL36, which is expressed early in infection and at reactivation but absent during latency (28, 31), may relieve RBPJ-dependent IE repression to facilitate lytic reactivation, although this remains to be examined. This work shows that miR-UL36 mirrors other HCMV miRNAs that target multiple nodes within a pathway (41, 42) highlighting the integral role that viral miRNAs play in regulating virus replication.

Latent infection of HPCs is a critical stage of the virus lifecyle. Here we show that disruption of UL8 endocytic motifs prevents Notch3 degradation and virus reactivation without affecting viral genome maintenance (Fig. 5C, S3A). Similarly, loss of miR-UL36 increased Notch3 and RBPJ levels, elevates HES1 expression, and blocks reactivation in HPCs (Fig. 6C–D). The Notch inhibitor CB-103 restores reactivation for both mutants, underscoring that suppression of Notch signaling is their central function. Furthermore, both mutants fail to reactivate in humanized mice (Fig. 7), confirming their essential roles in HCMV replication.

Collectively, HCMV employs two complementary mechanisms to inhibit Notch signaling in HPCs: UL8-mediated receptor degradation and miR-UL36–mediated post-transcriptional downregulation of pathway components. This dual strategy enforces a Wnt-ON/Notch-OFF signaling environment that promotes differentiation and reactivation. Given Wnt-Notch cross-talk, viral suppression of Notch may further potentiate Wnt activation, establishing a coordinated reactivation state. Intriguingly, each mutant fails to reactivate despite expression of the other regulator, indicating that UL8 and miR-UL36 function non-redundantly. The cooperation between UL8 and miR-UL36 parallels other viral regulatory pairs, such as UL7 with miR-US5-1 and miR-UL112, which act together to suppress FOXO3 (43). These convergent mechanisms underscore HCMV’s evolutionary strategy of pairing viral proteins and miRNAs to ensure robust control over host signaling.

While our study establishes UL8 and miR-UL36 as key Notch regulators, questions remain: How do they cooperate mechanistically to dismantle Notch signaling? What additional host transcripts are miR-UL36 targets, and do other viral miRNAs contribute? Could UL8 affect other Notch receptors or components? Addressing these questions will further illuminate how HCMV manipulates progenitor signaling to govern latency and reactivation.

## Materials and Methods

### Cells and media

Feeder-free hESCs were obtained from WiCell (WA01-H1, hPCSC Reg identifier (ID) WA001-A, NIH approval no. NIHhESC-10-0043). Cells were thawed and plated on Matrigel-coated six-well plates in complete mTeSR1 (Stem Cell Technologies). CD34^+^ HPCs were differentiated using a commercial feeder-free hematopoietic differentiation kit (STEMdiff Hematopoietic Kit, Stem Cell Technologies). HEK293 and adult normal human dermal fibroblasts (NHDF) were obtained from ATCC and cultured in Dulbecco’s modified Eagle’s medium (DMEM) supplemented with 10% heat-inactivated fetal bovine serum (FBS; Hyclone), 100 units/ml penicillin, 100 μg/ml streptomycin, and 100 μg/ml glutamine (Thermofisher). M2-10B4 and S1/S1 stromal cells were obtained from Stem Cell Technologies and maintained in DMEM with 10% FBS and penicillin, streptomycin, and glutamine as previously described (44). All cells were maintained at 37°C and 5% CO_2_.

### Reagents

siGENOME RISC-free control siRNA (Neg; Dharmacon), Notch3 siRNA (s453556; Thermofisher), and RBPJ siRNA (s145252, Thermofisher) were used in transfection experiments. Double stranded miRNA mimics were custom designed and synthesized by Integrated DNA Technologies. The following commercial antibodies were used: Notch3 (2889, Cell Signaling), RBPJ (5442, Cell Signaling), GAPDH (ab8245, Abcam), HCMV IE2 (MAB810, Sigma Aldrich), UL44 (Virusys Corporation, P1202-2), Turbo ID (AS204440, Agrisera). CB-103 was purchased from Selleckchem and the NanoGlo HiBiT Blotting System from Promega.

### Plasmids

The 4xCSL-luciferase reporter was constructed from the CBF1/pGL2-GLO TATA CAT plasmid, a gift from Dr. S. Speck. A fragment containing the multimerized high affinity CSL sites (4× CGTGGGAA) was excised by BamHI digest and ligated into a BglII/BamHI-digested RSV-TATA pGL2 vector, a gift from Dr. D. Towler. This modified vector has a TATA box from Rous sarcoma virus inserted into the pGL2-basic vector (Promega) to reduce the basal luciferase activity (Addgene plasmid #41726). Construct was sequenced for verification. hNICD3(3xFLAG)-pCDF1-MCS2-EF1-copGFP was a gift from Brenda Lilly (Addgene plasmid #40640). UL8 from TB40/E (EF999921) was modified to generate mutants in which tyrosine at position 305 was substituted with alanine, tyrosine at position 314 was substituted with alanine, or both tyrosines at positions 305 and 314 were substituted with alanine. These constructs were synthesized and cloned into pcDNA3.1 at GenScript. The Ad-HiBiT-UL8-Y305A-V5, Ad-HiBiT-UL8-Y314A-V5, Ad-HiBiT-UL8-Y305/314A-V5, were generated by subcloning the inserts from pCDNA3.1 into pGEM-T easy vector system and then into pAdTrack as previously described (21). Likewise, Ad-miR-67 and Ad-miR-UL36 were generated by subcloning 250bp up-and downstream of the sequences in their native context into the pAdTrack vector.

### Viruses

Viruses used in this study include BAC-generated WT TB40/E expressing GFP from the SV40 promoter. TB40/E mutant viruses containing point mutations in the pre-miRNA sequences for miR-UL36 were generated by galactokinase (galK)-mediated recombination (90). Briefly, the galK gene was inserted into the region of the pre-miRNA hairpin using homologous recombination (miR-UL36 galK F: GAAATAAGAAAAATCCACGCACGTTGAAAACACCTGGAAAGAACGTGCCCGAGCGAACGTCCTCTTTCCAGGTGTCCCTGTTGACAATTAATCATCGGCA, miR-UL36 galK R: GCTCCGTTCGCGCAACGCCCTGGGGCCCTTCGTGGGCAAGATGGGCACCGTCTGTTCGCAAGGTAAGCCCCACGCTCAGCAAAAGTTCGATTTATTCAAC. In the second recombination step, galK is removed using oligos that encompass the pre-miRNA sequence containing point mutations in the hairpin (miR-UL36 F: CACCTGGAAAGAACGTGCCCGAGCGAACGTCCTCTTTCCAGGTGTCAAGTTGctCGTGGGGCTTACCTTGCGAACAGACGGTGCCCATCTTGCCCACGAA, miR-UL36 R: TTCGTGGGCAAGATGGGCACCGTCTGTTCGCAAGGTAAGCCCCACGAGCAACTTGACACCTGGAAAGAGGACGTTCGCTCGGGCACGTTCTTTCCAGGTG). To generate the ΔmiR-UL36 FLAG and UL36 FLAG viruses, the following approach was taken. The galK gene was inserted into either WT TB40/E or the ΔmiR-UL36 BAC immediately upstream of the UL36 stop codon using the following primers: (UL36 FLAG galK F: TAAAATGCTGTATTATATACAAAAACATGCACATAGACAAACGGGACCACCGTGCTCGTCATCCCCTCCTTAATCACCTGTTGACAATTAATCATCGGCA, UL36 FLAG galK R: CGGTCGACGGCACATTATTCCTGGCGCCGCCAACGGCATGCCGCCCCTCACCCCGCCACACGCCTACATGAATAACCTCAGCAAAAGTTCGATTTA). The galactokinase gene was then replaced using oligos containing the FLAG epitope (UL36 FLAG oligo F: AAACGGGACCACCGTGCTCGTCATCCCCTCCTTAATCACTTGTCGTCATCGTCTTTGTAGTCGTTATTCATGTAGGCGTGTGGCGGGGTGAGGGGCGGCA and UL36 FLAG oligo R: TGCCGCCCCTCACCCCGCCACACGCCTACATGAATAACGACTACAAAGACGATGACGACAAGTGATTAAGGAGGGGATGACGAGCACGGTGGTCCCGTTT).

To generate a C-terminal fusion of the biotin ligase BirA to UL8 in the HCMV TB40/E strain, and to introduce two point mutations substituting tyrosine residues 305 and 315 with alanine, we employed a homologous recombination approach. Specifically, we used an adapted version of the en passant recombination method, which enables “scarless,” in-frame incorporation of genetic elements and site-directed mutations without leaving residual heterologous DNA sequences in the final viral construct (45, 46). Briefly, targeted alterations in the UL8 ORF were introduced using recombination primers carrying the desired nucleotide substitutions. Each primer contained a 50 bp sequence homologous to the target region in the HCMV genome, with an additional 50 bp of sequence either upstream (sense primer) or downstream (antisense primer) relative to the UL8 locus in the BAC. A 20 bp primer-binding site was added to the 3′ end of each primer to allow amplification of an aminoglycoside 3-phosphotransferase gene conferring kanamycin resistance (KanR), preceded by an I-SceI homing endonuclease recognition site (kanintoBirA-S: TATGTTTTGGCGCCTGAAGCGGGGACCAGCAGCAATCGGCCTGGGCCCGGTCATCGGAATTGTCATGGCAGAAGCGCTGCGAAAGCTGGGAGCAGACAAGTAGGGATAACAGGGTAATAAG; kanintoBirA-AS: GCCAGCTTTCTATCCTGCAGATACAGGTCATTGGGCCATTTGACTCGCACCTTGTCTGCTCCCAGCTTTCGCAGCGCTTCTGCCATGACAATTCCGATGAAGAGCGCTTTTGAAGCTGG. The resulting PCR fragment was recombined into the TB40/E BAC maintained in *E. coli* strain GS1783 by heat-shock induction of λ phage–derived Red recombination genes. Recombinant clones were selected on kanamycin-containing LB agar plates, cultured in LB medium, and verified by HindIII restriction digest and Sanger sequencing of the altered locus to confirm both genome integrity and correct cassette integration. To excise the KanR marker, I-SceI endonuclease was conditionally expressed from GS1783 by arabinose induction to generate a double-strand break at the recognition site, while Red recombination genes were simultaneously induced by heat shock. This allowed recombination across the duplicated 50 bp homology regions flanking the selection cassette. Correct loss of the antibiotic resistance marker was confirmed by differential growth on LB agar plates ± kanamycin, followed by HindIII restriction digest and Sanger sequencing of the altered locus. For fusion of the biotin ligase BirA to the C-terminus of UL8, we first introduced an I-SceI site followed by a KanR cassette, flanked by a duplicated 50 bp region located immediately upstream of the BirA integration site (RepairBirA-S: AGTTCCTCCGTCCTCTGCACCGACGGCGAAAACACCGTCGCGTCCGACGCAACGGTGACGGCATTAGCTAGCAAAGACAATACTGTGCCTCTGAAGCTGATAGGGATAACAGGGTAATAAG; RepairBirA-AS: TTCTCCCAGCTGTTCGCCACTATGGAACTCGCCATTAGCCAGGAGAGCGATCAGCTTCAGAGGCACAGTATTGTCTTTGCTAGCTAATGCCGTCACCGTTAGAGCGCTTTTGAAGCTGG). The BirA ORF, together with the KanR cassette flanked by 50 bp homology arms, was then PCR-amplified using primers that contained a 20 bp binding site at the 5′ and 3′ ends of BirA and an additional 50 bp of homology to the TB40/E BAC region immediately upstream and downstream of the UL8 termination codon (BirAontoUL8-S: CAGAACTGGGCAAGCCCATCCCCAACCCCCTGCTGGGCCTGGACAGCACCGCTAGCAAAGACAATACTGT; BirAontoUL8-AS: GTCATCCATAGAAATAACCGCAAACACTTCTTAGACATCATAGTATATTACTTTTCG GCAGACCGCAG). En passant recombination was performed as described above for UL8 Y305/314A point mutations (UL8 Y305/Y314A-S: ATCATTTTCATCATTTTTATTATCATCTGTCTACGAGCACCTCGAAAAGTTGCTGATCGTTGGAAAGACAGCAGAGAGGCCGGACAAGTGTTTATGACGGTAGGGATAACAGGGTAATAAG UL8 Y305/Y314A-AS: GCTGTCCAGGCCCAGCAGGGGGTTGGGGATGGGCTTGCCCAGTTCTGTGTCCGTCATAAACACTTGTCCGGCCTCTCTGCTGTCTTTCCAACGATCAGCAAGAGCGCTTTTGAAGCTGG), and selected clones were analyzed by restriction digest. All the final BAC clones were furthermore analyzed by next generation sequencing using an Illumina iSeq sequencer to exclude off-target mutations. All virus stocks were propagated and titered on NHDFs using standard techniques. To assess growth kinetics, NHDFs were infected at a MOI of 3 for single-step growth curves or a MOI of 0.01 for multi-step growth curves for 2 hr. Cell-associated and supernatant virus was harvested at multiple time points post-infection. Titers were determined by plaque assay on NHDFs. For recombinant adenovirus production, pAdTrack-CMV (Addgene plasmid #16405) and AdEasier-1 cells (Addgene #16399) were a gift from Bert Vogelstein. The pAdTrack plasmids were linearized by digesting with restriction endonuclease *Pme I*, and subsequently recombined into *E. coli* BJ5183 cells containing the adenoviral backbone plasmid pAdEasy-1 (AdEasier-1 cells). Recombinants were selected for kanamycin resistance, and the recombination confirmed by restriction endonuclease analyses. Finally, the recombinant plasmids were linearized with *PacI* before transfection with Lipofectamine 2000 (ThermoFisher) into the adenovirus packaging cell lines HEK293 cells. The control vector Ad-GFP, Ad-HiBiTUL7, Ad-HiBiTUL8-V5, Ad-HiBiTUL8-mutDTEL-V5, Ad-HiBiT-UL8-Y305A-V5, Ad-HiBiT-UL8-Y314A-V5, Ad-HiBiT-UL8-Y305/314A-V5, Ad-miR-67, Ad-miR-UL36 were produced, purified and titered in HEK293, as previously described (43).

For adenovirus transduction, hESC-derived CD34^+^ HPCs were resuspended at low volume in IMDM containing 10% BIT serum supplement, L-glutamine, low density lipoproteins, beta-mercaptoethanol, and stem cell cytokines as previously described (ref) in a low binding 24-well plate (Corning low-attachment Hydrocell). Cells were then infected with adenoviruses at an MOI of 125 for 4 hours with continual rocking then spin infected at 300g for 30min, resuspended, and cultured overnight. Culture conditions were supplemented with additional media and infection continued for a total of 48 hours. Samples were then FACS isolated for pure populations of transduced HPCs (viable, CD34^+^, GFP^+^) as previously described (43). NHDF were infected with adenoviruses at an MOI of 250 for 48 hours.

### Proximity biotinylation

10cm cell culture dishes containing 70–80% confluent monolayers of NHDF were infected at an MOI of 1 with either HCMV TB40/E-GFP-UL8-TurboID, or WT HCMV TB40/E-GFP. Three days post infection, cells were incubated for 6h in complete media supplemented with 50μg/mL biotin. Cells were lysed in Lysis buffer containing 50mM Tris-HCl, pH 8, 1% Nonidet P-40, 150 mM NaCl, and 1X protease inhibitor cocktail (P8340, Sigma), 1X phosphatase inhibitor cocktail 1 (P2850; Sigma), 1X phosphatase inhibitor cocktail 2 (P5726; Sigma) and insoluble debris were pelleted at 10,000xg for 10 min at 4°C. 1mg of protein lysates was incubated at 4°C overnight with 500 μl of streptavidin conjugated to magnetic beads (New England BioLabs, Ipswich, MA). Beads were washed once in 1.5 ml of wash buffer 1 (2% SDS in H_2_O), once with wash buffer 2 (0.1% deoxycholate, 1% Triton X-100, 500 mM NaCl, 1 mM EDTA, and 50 mM HEPES [pH 7.5]), once with wash buffer 3 (250 mM LiCl, 0.5% NP-40, 0.5% deoxycholate, 1 mM EDTA, 10 mM Tris [pH 8.1]), and then twice with wash buffer 4 (50 mM Tris [pH 7.4], 50 mM NaCl). To evaluate sample integrity, 25% of the total was retained for immunoblots. The remaining 75% of the sample (for analysis by mass spectrometry) was washed an additional two times in 50mM ammonium bicarbonate and then resuspended in 268μL 50mM ammonium bicarbonate and incubated on a 70°C heat block for 10 min with agitation. The samples were immediately treated with 132μL 6M urea and then cooled to room temperature before adding 2.5μL of fresh 0.5M TCEP (Tris(2-carboxyethyl) phosphine hydrochloride; Sigma) and incubated for 30min at room temperature, followed by adding 9μL of fresh 0.5M iodoacetamide and incubating in the dark for another 30 min at room temperature. The samples were then subjected to tryptic digestion by adding 3.7μL 10mM CaCl_2_ followed by 20μL of 0.1μg/μL sequencing grade trypsin and incubated overnight at 37°C with rotation. Twenty microliters of formic acid were then added to the eluate and stored at-80°C until LC-MS/MS analysis. Samples were desalted using ZipTip C18 (Millipore, Billerica, MA) and eluted with 70% acetonitrile/0.1% TFA (Trifluoracetic acid; Sigma) and the desalted material dried in a speed vac. On bead tryptic digests were analyzed by the Fred Hutchinson Proteomics Core Facility (Seattle, WA).

### Liquid chromatography tandem-mass spectrometry

Desalted samples were brought up in 2% acetonitrile in 0.1% formic acid (12μL) and 10μL of sample analyzed by LC/ESI MS/MS with a ThermoScientific Easy-nLC II nano HPLC system (Thermo Scientific, Waltham, MA) coupled to a tribrid Orbitrap Fusion mass spectrometer (Thermo Scientific, Waltham, MA). Peptide separations were performed on a reversed-phase column (75 μm × 400 mm) packed with Magic C18AQ (5-μm 100Å resin; Michrom Bioresources, Bruker, Billerica, MA) directly mounted on the electrospray ion source. A 90-minute gradient from 7% to 28% acetonitrile in 0.1% formic acid at a flow rate of 300nL/minute was used for chromatographic separations. The heated capillary temperature was set to 300°C and a static spray voltage of 2100 V was applied to the electrospray tip. The Orbitrap Fusion instrument was operated in the data-dependent mode, switching automatically between MS survey scans in the Orbitrap (AGC target value 500,000, resolution 120,000, and maximum injection time 50 milliseconds) with MS/MS spectra acquisition in the linear ion trap using quadrupole isolation. A 2 second cycle time was selected between master full scans in the Fourier-transform (FT) and the ions selected for fragmentation in the HCD cell by higher-energy collisional dissociation with a normalized collision energy of 27%. Selected ions were dynamically excluded for 30 seconds and exclusion mass by mass width +/-10 ppm.

Data analysis was performed using Proteome Discoverer 2.2 (Thermo Scientific, San Jose, CA). The data were searched against Uniprot Human and CRAPome (Mellacheruvu D, Wright Z, Couzens AL, Lambert JP, St-Denis NA, Li T, et al. The CRAPome: a contaminant repository for affinity purification-mass spectrometry data. Nat Methods. 2013;10(8):730–6. Epub 20130707. doi: 10.1038/nmeth.2557) data repositories (>25% cutoff). Trypsin was set as the enzyme with maximum missed cleavages set to 2. The precursor ion tolerance was set to 10 ppm and the fragment ion tolerance was set to 0.6 Da. Variable modifications included oxidation on methionine (+15.995 Da), carbamidomethyl on cysteine (+57.021 Da), and acetylation on protein N-terminus (+42.011 Da). Data were searched using Sequest HT (47). All search results were run through Percolator for scoring (48). Pathway enrichment analysis of the UL8-associated proteins was performed using gene set enrichment analysis (GSEA) against multiple pathway databases, including KEGG and Hallmark gene sets. Significantly enriched pathways were identified based on false discovery rate (FDR) values below 0.05.

### Immunoprecipitation

Lysates were prepared from transfected, adenoviral transduced or HCMV infected cells as described above. Soluble proteins were quantified with a Bicinchoninic acid (BCA) Assay (Thermo Fisher) and equal quantities of protein were used for immunoprecipitation. Immunoprecipitation was performed by incubating 300-500 μg of protein lysate with 1:100 of primary antibody for 3h at 4°C, followed by capture of antigen-antibody complexes with 20 μl of protein A or G magnetic beads (Cell Signaling). The resin beads were then rapidly washed five times with Tris-Buffered saline buffer with 0.1% Nonidet P-40. Immunoprecipitates were eluted from the affinity resin in 2X Laemmli Buffer for 10 min at 80°C and then subjected to SDS-Page and Western Blotting as described below.

### Luciferase assays

HEK293T cells were seeded into 96 well plates and transfected with 50 ng of 4XCSL-Luciferase (41726; Addgene) vector, 50ng 3XFLAGNICD3 (20185; Addgene) and 100 fmol of negative control or miRNA mimic using Lipofectamine 2000 (Invitrogen). Twenty-four hours after transfection cells were harvested for luciferase assay using the Dual-Glo Reporter Assay Kit (Promega) according to the manufacturer’s instructions. Luminescence was detected using a Veritas microplate luminometer (Turner Biosystems). All experiments were performed in triplicate and presented as mean +/-standard deviation.

### Western blot analysis

Cells were harvested in protein lysis buffer (50mM Tris-HCl pH 8.0, 150mM NaCl, 1% NP40, and protease inhibitors), loading buffer (4X Laemmli Sample Buffer with 2-mercaptoethanol) was added, and lysates were incubated at 95°C for 5 min. Extracts were loaded onto 4-15% acrylamide gels (Biorad), transferred to Immobilon-P membranes (Millipore), and visualized with the specified antibodies. The relative intensity of bands detected by Western blotting was quantified using ImageJ software.

### Quantitative RT-PCR

Reverse transcription-PCR (RT-PCR) was used to quantitate cellular transcripts in infected NHDFs or CD34^+^ HPCs. Total RNA was isolated from infected cells using Trizol. cDNA was prepared using 1000ng of total RNA and random hexamer primers for cellular and viral RNAs, and using 100ng total RNA and miRNA hairpin-specific primers for viral miRNAs. Samples were incubated at 16°C for 30 minutes, 42°C for 30 minutes, and 85°C for 5 minutes. Real-time PCR (Taqman) was used to analyze cDNA levels in infected samples. An ABI StepOnePlus Real Time PCR machine was used with the following program for 40 cycles: 95°C for 15 sec and 60°C for 1 minute. Primer and probe sets for HES1 (Hs00172878_m1), HEY1 (Hs05047713_s1) and 18S (Hs03928990_g1) were obtained from Thermo Fisher Scientific. Relative expression was determined using the ΔΔCt method using 18S as the standard control with error bars representing the standard deviation from at least 3 experiments.

### CD34^+^ HPC latency and reactivation assays

Differentiated hESCs were infected with the indicated viruses at an MOI of 2 for 48hr, or were left uninfected, in stem cell media (Iscove’s modified Dulbecco’s medium [IMDM] [Invitrogen] containing 10% BIT serum replacement [Stem Cell Technologies], penicillin/streptomycin, stem cell factor [SCF], FLT3 ligand [FLT3L], interleukin-3 [IL-3], interleukin-6 [IL-6] [all from PeproTech], 50uM 2-mercaptoethanol, and 20ng/ml low-density lipoproteins). Pure populations of viable, infected (GFP^+^) CD34^+^ HPCs were isolated by fluorescence-activated cell sorting (FACS) (BD FACSAria equipped with 488-, 633-, and 405-nm lasers and running FACSDiva software) and used in latency assays as previously described. Briefly, cells were cultured in transwells above irradiated stromal cells (M2-10B4 and S1/S1) for 12 days to establish latency. Virus was reactivated by coculture with NHDF in RPMI medium containing 20% FBS, 1% P/S/G, and 15ng/ml each of G-CSF and GM-CSF in an extreme limiting dilution assay (ELDA). GFP^+^ wells were scored 3 weeks postplating and the frequency of infectious centers was using ELDA software.

### Engraftment and infection of humanized mice

All animal studies were carried out in strict accordance with the recommendations of the American Association for Accreditation of Laboratory Animal Care. The protocol was approved by the Institutional Animal Care and Use Committee (protocol 0922) at Oregon Health and Science University. NOD-*scid*IL2Rγ_c_^null^ mice were maintained in a pathogen-free facility at Oregon Health and Science University in accordance with procedures approved by the Institutional Animal Care and Use Committee. Both sexes of animals were used. Humanized mice were generated as previously described. The animals (12-14 weeks post-engraftment) were treated with 1 ml of 4% Thioglycollate (Brewer’s Media, BD) by intraperitoneal (IP) injection to recruit monocyte/macrophages. At 24hr post-treatment, mice were infected with HCMV TB40/E-infected fibroblasts (approximately 10^5^ PFU of cell-associated virus per mouse) via IP injection. A control group of engrafted mice was mock infected using uninfected fibroblasts. Virus was reactivated as previously described (35).

### Quantitative PCR for viral genomes

DNA from CD34^+^ HPCs was extracted using the two-step TRIZOL (Thermofisher) method according to the manufacturer’s directions. Total DNA was analyzed in triplicate using TaqMan FastAdvanced PCR master mix (Applied Biosystems), and primer and probe for HCMV *UL141* and human β-globin as previously described (21). Copy number was quantified using a standard curve generated from purified HCMV BAC DNA and human β-globin-containing plasmid DNA, and data were normalized assuming two copies of β-globin per cell.

### Statistical analysis

Statistical analysis was performed using GraphPad Prism software (v10) for comparison between groups using student’s t-test, one-way or two-way analysis of variance (ANOVA) with Tukey’s post-hoc test or Bonferroni’s multiple comparison test as indicated. Values are expressed as mean +/-standard deviation or standard error of the mean, as indicated in the figure legends. Significance is highlighted with p<0.05.

## Acknowledgements

We thank Phil Gafken and Lisa Jones for help in carrying out MS experiments. We thank Andrew Townsend for graphics assistance. We gratefully acknowledge Jay Nelson, Daniel Streblow, Felicia Goodrum, and Andrew Yurochko for helpful discussions. This work was supported by grant P01 AI127335 from the National Institute of Allergy and Infectious Diseases, NIH funded to P.C. and M.H.H, and by National Institute of Health R37 AI21640 to M.H.H. R.T. is supported by the National Institute of Health under Award number T32AI170496. The funder had no role in study design, data collection and analysis, decision to publish, or preparation of the manuscript. This research utilized the Proteomics & Metabolomics Shared Resource of the Fred Hutch/University of Washington Cancer Consortium (P30 CA05704).

---

**Supplemental Figure 1: Treatment with CB-103 is not cytotoxic and does not impact lytic replication.** (A) CD34^+^ HPCs were incubated with 10 μM CB-103 or DMSO for 7 days. A colorimetric assay (WST-1 based, Roche) was used to determine cytotoxicity according to the manufacturer’s directions. Values are means±standard error (SD) (error bars). (B) NHDF cells were infected at an MOI of 0.05 and treated with 10 μM CB-103 or DMSO. Supernatants were harvested at the indicated times post-infection and titrated by TCID50.

**Supplemental Figure 2: Mutation of miR-UL36 does not affect pUL36 expression or virus replication.** (A) NHDF were infected with the indicated viruses at an MOI of 3. After 24 and 48 hours, protein lysates were harvested, and samples were immunoblotted using the indicated antibodies. (B) NHDF were infected with WT or ΔmiR-UL36 virus at an MOI of 3. At the indicated times post-infection, supernatant was harvested and titered on NHDF.

**Supplemental Figure 3: Characterization of viral mutants.** (A) Total DNA from primary CD34^+^ HPC cultures was extracted at 14 dpi and HCMV genomes were quantified using quantitative PCR with primers and probe specific for the *UL141* gene. Viral genomes were normalized to total cell number determined using human β-globin as a reference. Data shown is the mean value of three experiments and error bars represent standard deviation. (B) NHDF cells were infected with WT or UL8 Y305/314A mutant virus at an MOI of 0.05. Supernatants were harvested at the indicated times post-infection and titrated by TCID50. **(C)** Genome copy number as determined in (A). **(D)** NHDF cells were infected with WT or ΔmiR-UL36 mutant at an MOI of 0.05. Supernatants were harvested at the indicated times post-infection and titrated by TCID50.

